# The controlled direct effect of temperament at 2-3 years on cognitive and academic outcomes at 6-7 years

**DOI:** 10.1101/410506

**Authors:** Shiau Y. Chong, Catherine R. Chittleborough, Tess Gregory, John Lynch, Murthy N. Mittinty, Lisa G. Smithers

## Abstract

There is widespread interest in temperament and its impact upon cognitive and academic outcomes. Parents adjust their parenting according to their child’s temperament, however, previous studies have not accounted for parenting while estimating the association between temperament and academic outcomes. We examined the controlled direct effect of temperament (2-3 years) on cognitive and academic outcomes (6-7 years) when mediation by parenting practices (4-5 years) was held constant. Participants were from the Longitudinal Study of Australian Children (n=5107). Cognitive abilities were measured by the Peabody Picture Vocabulary Test (verbal) and the Matrix Reasoning test (non-verbal). Literacy and numeracy were reported by teachers using the Academic Rating Scale. Mothers reported children’s temperament using the Short Temperament Scale for Toddlers (subscales: reactivity, approach, and persistence). Parenting practices included items about engagement in activities with children. Marginal structural models with inverse probability of treatment weights were used to estimate the controlled direct effect of temperament, when setting parenting to the mean. All temperament subscales were associated with cognitive abilities, with persistence showing the largest controlled direct effect on verbal (β=0.58; 95%CI 0.27, 0.89) and non-verbal (β=0.19; 0.02, 0.34) abilities. Higher persistence was associated with better literacy (β=0.08; 0.03, 0.13) and numeracy (β=0.08; 0.03, 0.13), and higher reactivity with lower literacy (β=−0.08; −0.11, −0.05) and numeracy (β=−0.07; −0.10, −0.04). There was little evidence that temperamental approach influenced literacy or numeracy. Overall, there was a small controlled direct effect of temperament on cognitive and academic outcomes after accounting for parenting and confounders.

## Introduction

There is widespread interest in whether children’s temperament influences their cognitive and academic outcomes [1, 2]. Temperament is the individual characteristics in behavioral styles that are biologically-based, but also shaped by experiences and environment [3]. Three aspects of temperament thought to impact learning and cognition are reactivity, persistence and approach [4]. Reactivity encompasses a child’s emotional intensity and volatility [4], and may interfere with the child’s learning processes [1, 5]. For example, a highly reactive child may become easily frustrated and find it difficult to learn [6]. Children with high persistence are more likely to stay on task and maintain their attention despite distractions [4], which has benefits for learning [7]. Temperamental approach is the degree of comfort experienced when encountering new situations or people [4]. Low approach may present challenges for young children in the transition to school as they are faced with many new situations, teachers and peers [8]. Temperamental reactivity and persistence are modifiable [9, 10]. For instance, a cluster-randomized trial showed that an intervention to develop children’s persistence, attention, and impulse control resulted in improvements to academic outcomes [10]. Increasing children’s persistence and reducing their reactivity may be a mechanism to improve children’s cognitive and academic outcomes, provided that there are direct effects of temperament on these outcomes [9-11].

To estimate the direct effect of temperament, we need to account for the fact that children’s temperament may influence parenting [12, 13], which in turn, is known to influence children’s cognitive and academic outcomes [14]. Maccoby *et al* [13] showed that temperamentally difficult children (*i.e.* high emotional intensity, difficult to calm) received less teaching from parents at 18 months than temperamentally easy children. However, Dixon *et al* [12] found mothers engaged in more high quality play with temperamentally difficult than easy children. Parental engagement in play and cognitive stimulation activities has positive impacts on children’s outcomes [14], and differential parental engagement for temperamentally easy and difficult children might, in part, explain effects of temperament on cognitive and academic outcomes.

While some studies have examined the direct effect of temperament on cognitive and academic outcomes, most involve limited adjustment for confounding [2, 5, 15]. The few studies that accounted for parenting suggest a direct effect of temperament on outcome [16, 17]. However, simple adjustment for parenting practices could introduce bias when parenting practices are affected by temperament (parenting is a mediator) and when there are confounders of parenting and outcomes (mediator-outcome confounding; Supplementary File S1) [18]. In the current study, we use traditional linear regression models to examine the total effect of temperament (reactivity, approach, persistence) at 2-3 years on cognitive and academic outcomes at 6-7 years, simply to compare results with past studies. In our main findings, we use marginal structural models (MSMs) and counterfactual theory to estimate the controlled direct effects (CDEs) of temperament on cognitive and academic outcomes, while accounting for parenting practices at 4-5 years. MSMs adjust for confounding using inverse probability of treatment weighting [19]. MSMs allow control of parenting by setting it to some uniform value, which in turn, enables the estimation of the ‘controlled’ direct effect of temperament on outcomes. Here parenting is somewhat of a nuisance variable, we need to account for it but it is not part of the effect we want to estimate. Another advantage of MSMs over traditional methods is weighting for mediator-outcome confounding, as this does not involve statistical adjustment for the intermediate variable, which could introduce bias (Supplementary Material S1) [20].

## Methods

### Study design and sample

Data were from the Longitudinal Study of Australian Children (LSAC). LSAC is a population-based study that commenced in 2004. Participants were recruited using a two-stage clustered sampling process [21]. At commencement, 5107 infants (mean age 8.8 months) were recruited and followed-up at 2-3 (n=4606), 4-5 (n=4386), and 6-7 (n=4242) years. LSAC is considered broadly representative of Australian children [21].

### Ethical approval and consent

All procedures performed in studies involving human participants were in accordance with the ethical standards of the institutional and/or national research committee and with the 1964 Helsinki declaration and its later amendments or comparable ethical standards. LSAC was approved by the Australian Institute of Family Studies ethics committee. Written informed consent was obtained from all participants’ caregivers.

### Data

De-identified data used in the current study from LSAC is accessible to *bona fide* researchers by application. Further information is available at the ‘Accessing LSAC data’ webpage (https://growingupinaustralia.gov.au/data-and-documentation/accessing-lsac-data).

### Cognitive ability and academic achievement (*Y*)

Verbal ability (receptive vocabulary) was measured using an adapted Peabody Picture Vocabulary Test (PPVT). The adapted PPVT-III [22] was administered by a trained interviewer to children aged 6-7 years during home interviews. The child pointed to the picture that best represented the meaning of a word spoken by the examiner [23]. The adapted PPVT-III was comparable to the full PPVT-III (correlations ranging 0.93–0.97) with high internal consistency (person-separation reliability 0.76) [22]. Scale scores were created using Rasch modelling (Mean=64, SD=8) [22].

Non-verbal ability (fluid reasoning) was measured using the Matrix Reasoning test from the Wechsler Intelligence Scale for Children, 4^th^ edition [24]. The Matrix Reasoning test comprised 35 items. The child was presented with an incomplete set of diagrams and asked to select the picture that completes the set from 5 different options. Scores were reported as standard scores, from age-appropriate norms (mean=10, SD=3). High internal consistency of the Matrix Reasoning test has been established in normative samples of Australian children (Cronbach’s α=0.88 for 6-year-olds, 0.91 for 7-year-olds) [24].

Academic achievement at 6-7 years was measured using the adapted Academic Rating Scale (ARS) [25], which has two subscales: literacy (10 items) and numeracy (9 items). Teachers rated the child’s skills and knowledge in relation to other children of the same age from ‘not yet demonstrated skill’ to ‘demonstrates skill competently and consistently’. Literacy items included ‘reads books fluently’ and ‘writes sentences with more than one clause’. Numeracy items included ‘uses a variety of strategies to solve math problems’ and ‘makes reasonable estimates of quantities’. Total scores ranging from 1 to 5 were created using Rasch modelling. Higher scores indicate higher proficiency. The ARS has high internal reliability (Cronbach’s α=0.96 for literacy, 0.95 for numeracy) [26].

### Temperament (*X*)

Temperament was measured at 2-3 years using the Short Temperament Scale for Toddlers (STST) [27]. The STST was adapted from the Toddler Temperament Scale [28]. The STST consists of 3 subscales (4 items each): reactivity, approach, and persistence, rated by the primary caregiver (98.2% mothers) from 1 (almost never) to 6 (almost always). Average scores were calculated for each subscale. Higher scores indicate higher reactivity (more negative emotion), higher approach (lower shyness), and higher persistence. All subscales had high internal reliability in the current sample (α=0.99 for each subscale).

### Parenting (*M*)

Parenting was assessed at 4-5 years using the home activities index which contains 7 items measuring how often mothers read to the child, tell stories, draw pictures, play indoor games, outdoor games, music, and involve the child in activities such as cooking or pet care. These items have been used as indicators of the quality of home environment, such as frequency of parent-child activities in UNICEF surveys [29]. Items were rated from 0 (none) to 3 (everyday), and a total score (0-21) derived by summing item scores. Higher scores indicates more positive parenting practices.

### Confounders of the association between temperament and outcomes (*C*)

Factors that might confound the associations between temperament and cognitive and academic outcomes were decided *a priori* using a directed acyclic graph (Supplementary Material S1). Confounders included indicators of socioeconomic position, intrauterine, child, maternal and family factors. These confounders were reported by mothers when children were 0-1 year. Details about how these confounders were measured are included in Supplementary Material S2.

### Confounders of the association between parenting practices and outcomes (*L*)

To estimate the CDEs, we need to account for confounding associated with parenting and cognitive or academic outcomes [18]. This set of confounders were reported by mothers at ages 4-5 years and included variables that were affected by temperament and in turn confound the parenting-outcome association (maternal psychological distress, number of siblings, maternal working status, household income, and financial hardship).

### Analysis

Model 1 estimates the total effect of temperamental reactivity, approach or persistence (*X*) on cognitive and academic outcomes (*Y*) after adjusting for confounders (*C*) using linear regression. Model 2 includes further adjustment for mediators (*M*) and confounders (*C, L*). Most studies have used traditional regression models 1 and 2, and these are provided for comparison with previous work. However, our *a priori* primary analysis (model 3) involved MSM. Using MSMs, we estimated the CDEs of temperament (reactivity, approach, or persistence, *X*), on cognitive and academic outcomes (*Y*) after accounting for parenting practices *(M),* potential confounders of the association between temperament and cognitive and academic outcomes *(C)*, and confounders of parenting practices and outcomes (*L*). Unlike traditional regressions, which estimate the conditional effect, the MSM founded on counterfactual theory estimates the marginal effect of temperament on cognitive and academic outcomes. The marginal effect is the difference in outcomes under the observed (*x*) and counterfactual exposures (*x*\*) of each individual. The mediator *M* (parenting) is set to a uniform level of *m* such that effects of temperament on outcomes are not mediated by parenting.[18] We used the log-likelihood ratio to test for interactions between temperament and parenting. As no interactions were found, the CDE generates the same result regardless of the level at which the mediator is set. Therefore, we set the mediator to its mean value (*m*). The CDEs were estimated from linear regression models of the form:

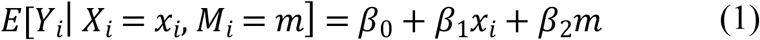

Potential confounding was accounted by fitting the model above with stabilized inverse probability weights of the form 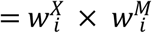, where

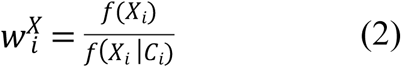

and

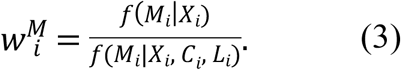

The weight 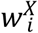 accounted for the confounding of the association between temperament and cognitive and academic outcomes by conditioning on *C*. The weight for the mediator 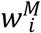 accounted for confounding of the association between parenting and cognitive and academic outcomes by conditioning on *X, C*, and *L*. Since the mediator was a continuous variable, probabilities were taken from the density functions. For computing the probabilities we assume that the mediator is normally distributed and hence used the normal density function. In the normal density function, parameters used to obtain probabilities were the observed mediator value, the predicted means and the root mean square error, estimated from linear regression [18]. Weights were truncated at the 1^st^ and 99^th^ percentile to deal with outliers. MSMs were performed separately for temperament reactivity, approach, and persistence. We performed sensitivity analysis to determine the extent to which an unmeasured confounder *U* might affect the association between temperament and cognitive and academic outcomes. We estimated the bias for the CDEs under conditions varying in prevalence and effect size of *U* (Supplementary Material S3) [30].

Analyses were performed using STATA 13.0 (StataCorp, College Station, TX).

### Multiple imputation

Twenty imputed datasets were generated under the missing at random assumption [31]. The imputation model included temperament, parenting practices, cognitive and academic outcomes, confounding variables and auxiliary variables that predicted missingness (parenting self-efficacy, temperament sociability, persistence, and reactivity at 4-5 years). We also performed analyses on the sample with observed outcome and the results were similar to the imputed sample (data not shown). Results from the imputed sample (n=5107) are reported.

## Results

Table 1 shows the characteristics of LSAC response, complete case (*n*=1647) and imputed (*n*=5107) samples. The highest proportion of missing data was teacher-reported outcomes of literacy and numeracy, although data from over 3300 children were available for these outcomes. Characteristics of the imputed sample were similar to the response sample.

**Table 1.** Characteristics of response, complete case, and imputed samples

|  | **Response sample<sup>a</sup>** |  | **Complete case**<br>**sample,<sup>b</sup>**<br>**n=1647** | **Imputed**<br>**sample,**<br>**n=5107** |
| --- | --- | --- | --- | --- |
|  | **N** | **M (SD) or %** | **M (SD) or %** | **M (SD) or %** |
| **Outcomes, *Y* (Cognitive ability and academic achievement)** |  |  |  |  |
| PPVT | 4185 | 74.4 (5.2) | 75.2 (4.9) | 74.2 (5.2) |
| Matrix reasoning | 4180 | 10.7 (3.0) | 11.0 (3.0) | 10.7 (3.0) |
| ARS-Literacy | 3408 | 3.4 (0.8) | 3.5 (0.7) | 3.4 (0.8) |
| ARS-Numeracy | 3357 | 3.3 (0.8) | 3.5 (0.8) | 3.3 (0.8) |
| **Exposures, *X* (Temperament)** |  |  |  |  |
| Reactivity | 3530 | 3.0 (1.0) | 2.9 (0.9) | 3.0 (0.9) |
| Approach | 3533 | 3.9 (1.0) | 4.0 (1.0) | 3.9 (1.0) |
| Persistence | 3532 | 4.3 (0.7) | 4.3 (0.7) | 4.3 (0.7) |
| **Intermediate variable, *M*** |  |  |  |  |
| Parenting practices | 4385 | 11.8 (3.9) | 12.0 (3.8) | 11.7 (3.9) |
| **Confounders of *X-M* or *X-Y*** |  |  |  |  |
| Mother's highest education, % |  |  |  |  |
| Tertiary | 1677 | 32.9 | 41.4 | 32.9 |
| Diploma/certificate | 1766 | 34.6 | 34.2 | 34.6 |
| Schooling only | 1656 | 32.5 | 24.4 | 32.5 |
| Hardship score | 5089 | 0.9 (1.3) | 0.7 (1.1) | 0.9 (1.3) |
| Housing tenure, rented or other, % | 5100 | 35.8 | 23.0 | 35.8 |
| Aboriginal or Torres Straits Islander, % | 5107 | 4.5 | 2.1 | 4.5 |
| Index of relative socio-economic disadvantage (IRSD) | 5107 | 1008.8 (60.2) | 1017.2 (57.2) | 1008.8 (60.2) |
| Child is male, % | 5107 | 51.1 | 48.5 | 51.1 |
| Birthweight for gestational age z-score | 4999 | 0.0 (1.1) | 0.1 (1.0) | 0.0 (1.1) |
| Duration of breast feeding, % |  |  |  |  |
| Never breastfed | 420 | 9.2 | 5.8 | 9.2 |
| <1 month | 518 | 11.4 | 8.4 | 11.4 |
| <3 months | 473 | 10.4 | 8.7 | 10.3 |
| <6 months | 692 | 15.2 | 14.0 | 15.2 |
| 6 months or more | 2461 | 53.9 | 63.2 | 54.0 |
| Maternal age | 5106 | 31.0 (5.5) | 32.2 (4.6) | 31.0 (5.5) |
| Mother is born in Australia, % | 4997 | 80.0 | 83.0 | 79.4 |
| Mother's psychological distress | 4308 | 4.4 (0.8) | 4.5 (0.5) | 4.4 (0.6) |
| Mother and partner argumentative relationship | 3931 | 2.2 (0.6) | 2.1 (0.6) | 2.2 (0.6) |
| Single-parent household, % | 5103 | 9.7 | 0.2 | 9.7 |
| Mother had gestational hypertension, % | 4238 | 7.8 | 7.8 | 8.1 |
| Mother had gestational diabetes, % | 4223 | 5.7 | 5.2 | 5.9 |
| Any alcohol during pregnancy, % | 4227 | 38.9 | 44.1 | 37.3 |
| Smoking during pregnancy, % | 4239 | 16.7 | 10.6 | 18.2 |
| **Confounders of *M-Y*** |  |  |  |  |
| Mother's psychological distress | 3818 | 4.5 (0.6) | 4.5 (0.5) | 4.5 (0.6) |
| Number of siblings | 4386 | 1.5 (1.3) | 1.5 (0.9) | 1.5 (1.0) |
| Mother's working status, % |  |  |  |  |
| Working full-time | 957 | 21.8 | 19.8 | 21.1 |
| Working part-time | 1804 | 41.2 | 46.6 | 40.7 |
| Currently not working | 1622 | 37.0 | 33.6 | 38.2 |
| Household income per week, \$ | 3668 | 1977.8<br>(1343.4) | 2054.5<br>(1365.3) | 1893.6<br>(1297.7) |
| Hardship score | 4365 | 0.3 (0.7) | 0.2 (0.6) | 0.3 (0.8) |

PPVT, Peabody Picture Vocabulary Test; ARS, Academic Rating Scale; *X-M*, exposure-
mediator; *X-Y*, exposure-outcome; *M-Y*, mediator-outcome
<sup>a</sup> Response sample is the number of participants who responded to specific assessment for each child exposure, outcome, or confounder,
<sup>b</sup> Complete case sample includes participants with complete data on all exposures, outcomes, and confounders.

Table 2 displays the effect estimates of temperament subscales on child outcomes using regression models adjusted for *C* (Model 1), *C, M*, and *L* (Model 2) and MSM (Model 3). The CDEs estimated from the MSM (Model 3) were closer to total effects (Model 1), while conventional regression (Model 2) underestimated effects. The MSM showed higher reactivity had negative effects on all outcomes, particularly verbal ability (β=−0.37 95% CI − 0.59, −0.14). Higher approach had positive effects on verbal and non-verbal abilities but little or no effect on literacy or numeracy. Higher persistence had positive impacts on all outcomes. Among the four outcomes, the largest effects were for verbal ability but effect sizes were small. In the MSM for instance, 1-unit higher persistence (range 1-5) was associated with 0.58-unit (0.11 SD) higher verbal and 0.19-unit (0.06 SD) higher non-verbal ability.

**Table 2:** Effect estimates of temperament reactivity, approach, and persistence at ages 2 to 3 years on child outcomes at ages 6 to 7 years (n=5107)

|  | **Model 1,**<br>**total effect** |  | **Model 2,**<br>**conventional regression** |  | **Model 3,**<br>**controlled direct effect** |  |
| --- | --- | --- | --- | --- | --- | --- |
| | $\beta$ | (95% CI) | $\beta$ | (95% CI) | $\beta$ | (95% CI) |
| **Reactivity** |  |  |  |  |  |  |
| PPVT | -0.38 | (-0.59, -0.17) | -0.36 | (-0.56, -0.15) | -0.37 | (-0.59, -0.14) |
| Matrix reasoning | -0.09 | (-0.18, 0.01) | -0.09 | (-0.19, 0.002) | -0.11 | (-0.21, -0.01) |
| ARS-Literacy | -0.07 | (-0.10, -0.04) | -0.07 | (-0.10, -0.04) | -0.08 | (-0.11, -0.05) |
| ARS-Numeracy | -0.06 | (-0.09, -0.03) | -0.06 | (-0.09, -0.03) | -0.07 | (-0.10, -0.04) |
| **Approach** |  |  |  |  |  |  |
| PPVT | 0.46 | (0.28, 0.65) | 0.40 | (0.21, 0.59) | 0.45 | (0.22, 0.67) |
| Matrix reasoning | 0.10 | (0.01, 0.19) | 0.08 | (-0.01, 0.18) | 0.11 | (0.01, 0.21) |
| ARS-Literacy | 0.03 | (-0.01, 0.06) | 0.02 | (-0.01, 0.06) | 0.03 | (-0.01, 0.07) |
| ARS-Numeracy | 0.02 | (-0.02, 0.05) | 0.01 | (-0.03, 0.05) | 0.01 | (-0.02, 0.05) |
| **Persistence** |  |  |  |  |  |  |
| PPVT | 0.61 | (0.38, 0.84) | 0.55 | (0.32, 0.79) | 0.58 | (0.27, 0.89) |
| Matrix reasoning | 0.16 | (0.04, 0.29) | 0.17 | (0.05, 0.30) | 0.19 | (0.02, 0.34) |
| ARS-Literacy | 0.08 | (0.04, 0.12) | 0.08 | (0.04, 0.12) | 0.08 | (0.03, 0.13) |
| ARS-Numeracy | 0.08 | (0.04, 0.12) | 0.09 | (0.05, 0.13) | 0.08 | (0.03, 0.13) |

*Note.* Model 1 is adjusted for confounders, $C$ . Model 2 is the conventional regression model including $X$ , $M$ , and all confounders $C$ and $L$ . Model
3 is a marginal structural model including $M$ and weighted for all confounders $C$ and $L$ .
$\beta$ are unstandardized coefficients.

The sensitivity analysis showed that the CDEs were generally robust in the presence of a binary unmeasured confounder (Supplementary Material S3). The observed CDEs would be explained by an unmeasured confounder if its prevalence differed between the exposed (*x*) and counterfactual (*x\**) by ≥80% and the estimated mean of the outcome differed by ≥0.60 within the two levels of the unmeasured confounder.

## Discussion

We found evidence for a direct effect of temperament at 2-3 years on children’s cognitive abilities and academic outcomes at ages 6-7 years. Of the three temperament dimensions, persistence had the largest CDE with 1-unit increase in persistence (5-point Likert scale) associated with effects in the order of 0.11 SD for verbal ability, 0.10 SD for literacy and numeracy and 0.06 SD for non-verbal reasoning. This is consistent with studies measuring temperamental persistence using different questionnaires [1, 7], children of different ages and using different statistical approaches [1]. For instance, a cross-sectional study of effortful control measured using the Child Behavior Questionnaire (CBQ) at 3-5 years was associated with 0.29 SD higher letter knowledge and 0.17 SD math achievement [7]. Rudasill *et al* [15] reported attention at 4.5 years measured with the CBQ was associated with 0.18 SD higher reading scores and 0.14 SD mathematic scores in 8-10 year-olds.

Reactivity was negatively associated with cognitive and academic outcomes, with the largest effect of 0.10 SD on literacy. This was similar to studies of preschoolers [1, 5] where CBQ temperament scores at 5.6 years from parents and teachers were averaged and standardized effects of ∼0.18-0.28 SD on mathematics and reading were observed [5]. The similarity in effects reported across studies from different countries, at different ages, as well as the use of different tools for measuring temperament adds strength to these findings.

Consistent with previous work involving the STST [8], higher scores on approach (lower shyness), were associated with higher verbal (0.09 SD) and non-verbal (0.04 SD) cognitive abilities. While we found little evidence of temperament approach on literacy and numeracy, others have reported that shy children were more likely to have poorer academic achievement [32]. Differences in these findings may be due to the small (n=125) cross-sectional design or because shyness was measured at 9-13 years, when children were expected to have more developed sociability skills.

Few studies have accounted for parenting when investigating the direct effect of temperament on cognitive and academic outcomes, and only narrow aspects of parenting (e.g. involvement in schooling[16], joint attention [17]) have been examined. We defined parenting as the frequency parents engaged in activities with their children because there is evidence this is important for children’s development [14]. Broader definitions of parenting might reveal different findings. Given that the CDEs of temperament on cognitive and academic outcomes were small, future research could examine temperament within the context of parenting practices, for instance, the effect of parenting on cognitive and academic outcomes may be heightened in temperamentally difficult compared with easy children. Parenting interventions could specifically target children with difficult temperament if they were more susceptible to the impact of parenting on their cognitive and academic outcomes.

Strengths of this study include the use of a large nationally-representative sample, prospective follow-up and teacher-reports of cognitive and academic outcomes. Previous studies are limited by small, non-representative samples, cross-sectional or short-term design (1-2 years) [33]. For instance, Valiente et al studied emotionality and academic abilities 6-months later in 291 children [5]. Furthermore, our methods account for parenting which influences children’s cognitive and academic outcomes, and a wide range of potential confounders.

Study limitations may include the dimensions of temperament used, mothers as informants, and measurement of parenting. We were limited to using the STSC because it was the only temperament measure available in LSAC. Nevertheless, our findings are consistent with studies using different temperament tools. Although we expect mothers to have excellent knowledge of their child’s temperament, it has been suggested that reporting of temperament might differ among mothers from lower socioeconomic position or suffering depression [34]. Mothers also reported parenting practices according to time engaging the child in daily activities, rather than the quality of the interactions. Finally, although we accounted for parenting, there may be other pathways that temperament could influence cognitive and academic outcomes, such as peer and teacher relationships, child care and birth order.

There is evidence of a CDE of temperamental persistence, approach and reactivity at ages 2-3 years on cognitive and academic outcomes at ages 6-7 years, but the effects are small. The largest CDE observed was for persistence on verbal ability but corresponded to only a 0.11 SD increase in verbal ability for every 1 point increase in persistence.

## Acknowledgements

We gratefully acknowledge the use of data from Growing Up in Australia, the Longitudinal Study of Australian Children (LSAC). The study is conducted in partnership between the Department of Families, Housing, Community Services and Indigenous Affairs (FaHCSIA), the Australian Institute of Family Studies (AIFS) and the Australian Bureau of Statistics (ABS). The findings and views reported in this paper are the authors’ and should not be attributed to FaHCSIA, the AIFS, or the ABS.

All authors contributed to the idea for the study. SYC acquired data from the Longitudinal Study of Australian Children to conduct the analyses, all authors had access to the data. Statistical advice was provided by MNM, with input from all authors. All authors approved the final manuscript.

## Supporting Information

**S1 File. The marginal structural model**

**S2 File. Confounders of the association between temperament and outcomes**

**S3 File. Sensitivity analysis for controlled direct effects**

